# Identifying G protein-coupled receptor dimers from crystal packings

**DOI:** 10.1101/282178

**Authors:** Ronald E. Stenkamp

## Abstract

Dimers of G protein-coupled receptors are believed to be important for signaling with their associated G proteins. Low resolution electron microscopy shows rhodopsin dimers in native retinal membranes, and CXCR4 dimers are found in several different crystal structures. Evidence for dimers of other GPCRs is more indirect. An alternative to computational modeling studies is to search for parallel dimers in the packing environments of the reported crystal structures of GPCRs. Two major structural types of GPCR dimers exist (as predicted by others), but there is considerable structural variation within each cluster. The different structural variants described here might reflect different functional properties and should provide a range of model structures for computational and experimental examination.

**Synopsis:** Analysis of intermolecular interactions in G protein-coupled receptor crystal structures shows two major types of dimers.

## 1. Introduction

Do G protein-coupled receptor (GPCR) dimers exist, and if they do, what do they look like? These are long-standing questions important for understanding GPCR function. Images of ordered arrays of rhodopsin in native membranes (Fotiadis et al., 2003, Liang et al., 2003, Filipek et al., 2004, Fotiadis et al., 2006) are consistent with dimers and higher oligomers, and electron microscopic studies of transducin/rhodopsin complexes further indicate that rhodopsin dimers exist and are physiologically functional (Jastrzebska et al., 2011, Jastrzebska et al., 2013). Other GPCRs are also believed to form dimers (Palczewski & Orban, 2013, Ferre et al., 2014, Kasai & Kusumi, 2014, Vischer et al., 2015, Gahbauer & Böckmann, 2016, Farran, 2017, Tian et al., 2017).

Crystal structure determinations of GPCRs have shown their basic protein folding topology of seven transmembrane helices (TM1 through TM7), and in some GPCRs, an eighth helix (H8) oriented parallel to the membrane surface. Analysis of crystal packing interactions (Katritch et al., 2013) led to the suggestion of two possible interfaces involved in dimer formation. One such interface involves transmembrane helices TM1 and TM2 as well as helix H8. Another interface of interest makes use of TM3, TM4, TM5 and TM6. These interfaces are small in area, with no amino acid residues sufficiently buried to be considered “core” residues. Computational analysis of these interfaces (Duarte et al., 2013) suggests that most of them do not result in physiologically relevant dimers. More recently, Baltoumas, *et al*. (Baltoumas et al., 2016) used molecular dynamics calculations to investigate the stability and dynamics of potential dimers extracted from crystal structures along with dimers modeled de novo. Their results are consistent with these two major candidates for dimer interfaces.

Over 200 crystal structures of GPCRs have been deposited in the Protein Data Bank (PDB) (Berman et al., 2000) since the structure of rhodopsin was solved in 2000 (Palczewski et al., 2000). This project investigates whether there are examples of GPCR-GPCR interactions in crystals that can be models of functional GPCR dimers. These might provide starting points for computational modeling or other experimental approaches probing for physiological dimers of biomedical interest. Reported here are the results of looking for GPCR dimers in the crystal structures.

## 2. Materials and Methods

A list of GPCR crystal structures was assembled by collating the results of keyword searches at the PDB website and lists found in review articles (Okada, 2012, Piscitelli et al., 2015) and in GPCR database websites (GPCRdb, http://gpcrdb.org/ (Isberg et al., 2016); Membrane Proteins of Known 3D Structure, http://blanco.biomol.uci.edu/mpstruc/). Table 1 lists the 215 entries in the initial set of crystal structures and the 121 crystal forms containing GPCR transmembrane domains.

**Table 1.**
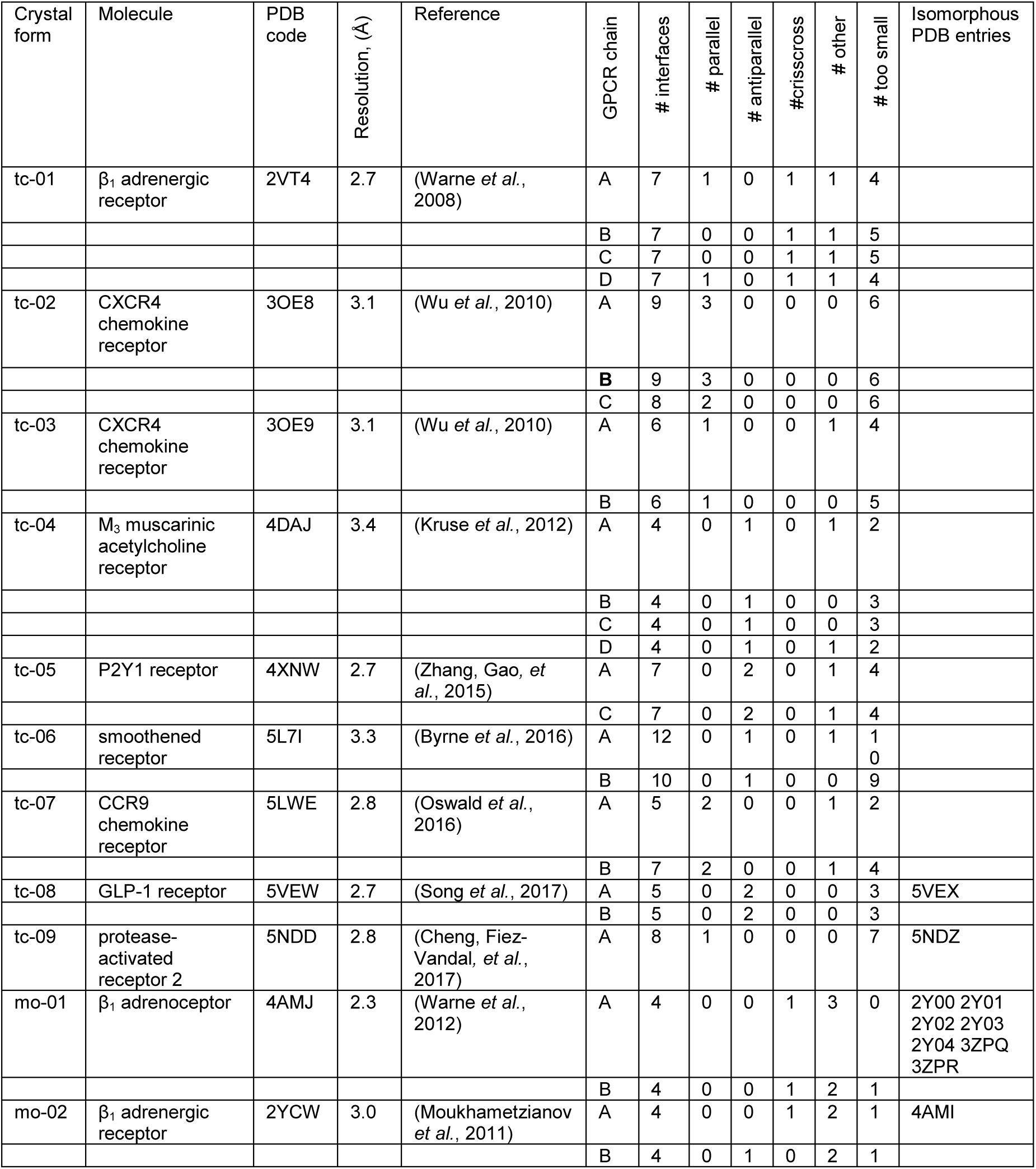
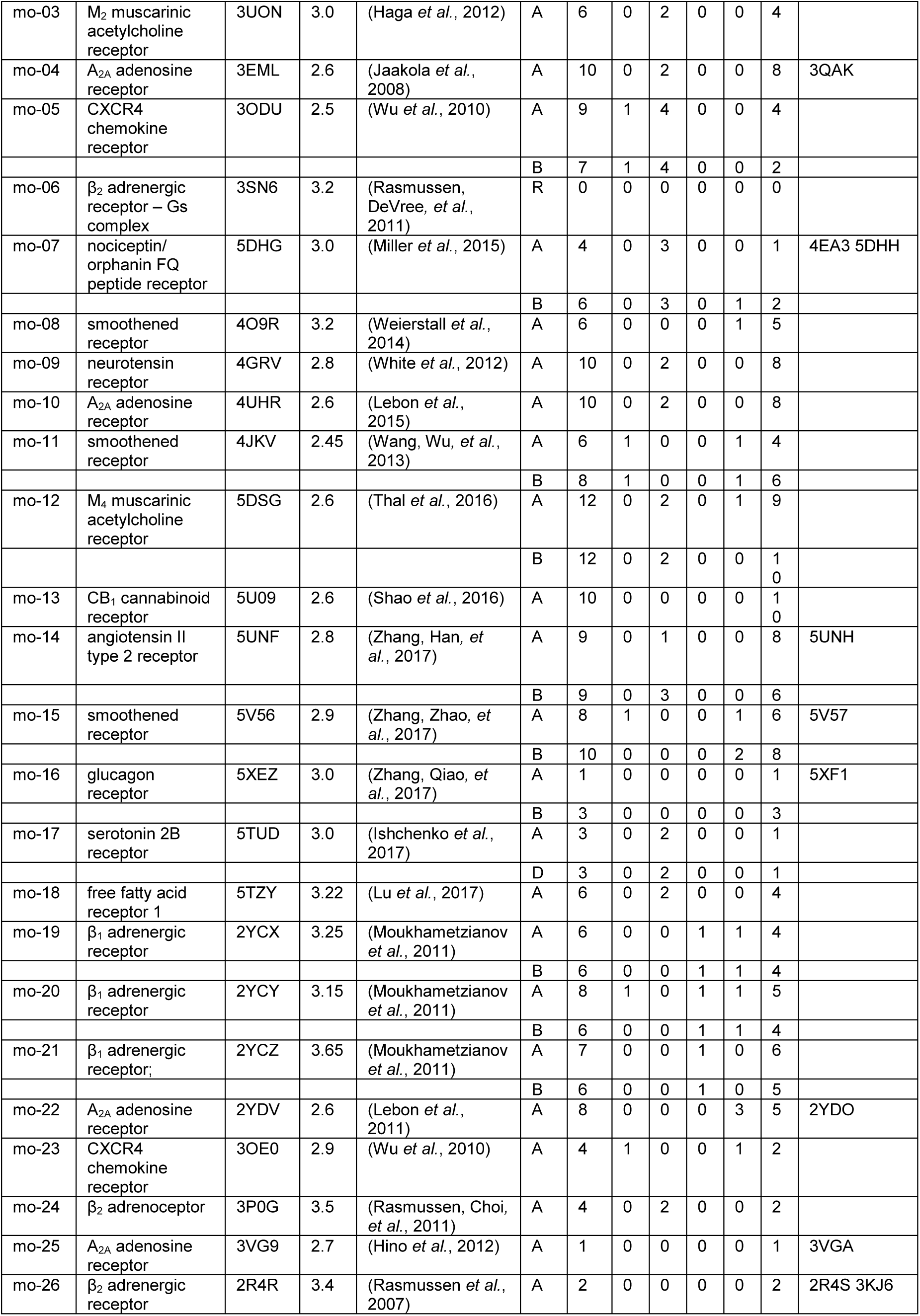
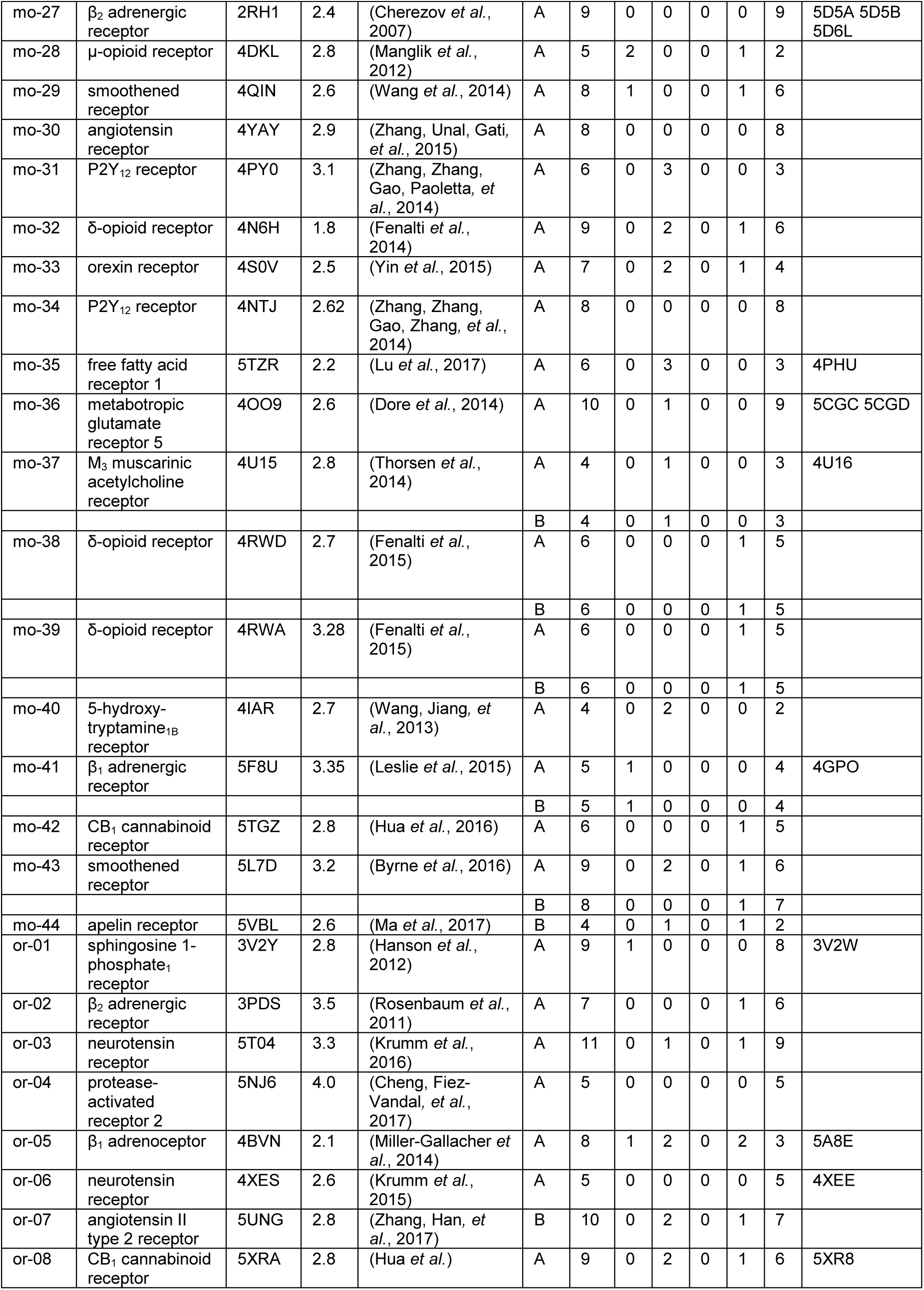
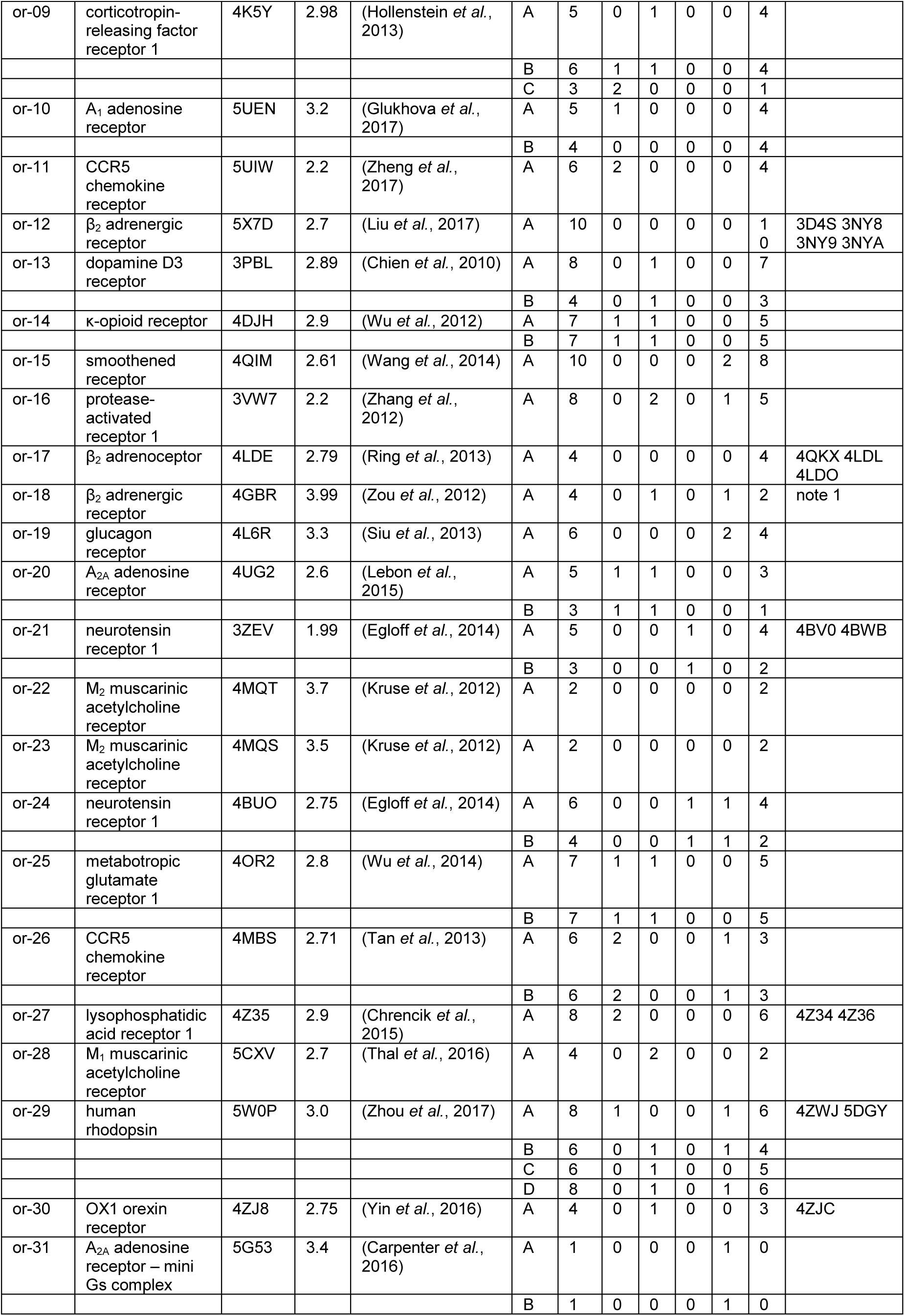
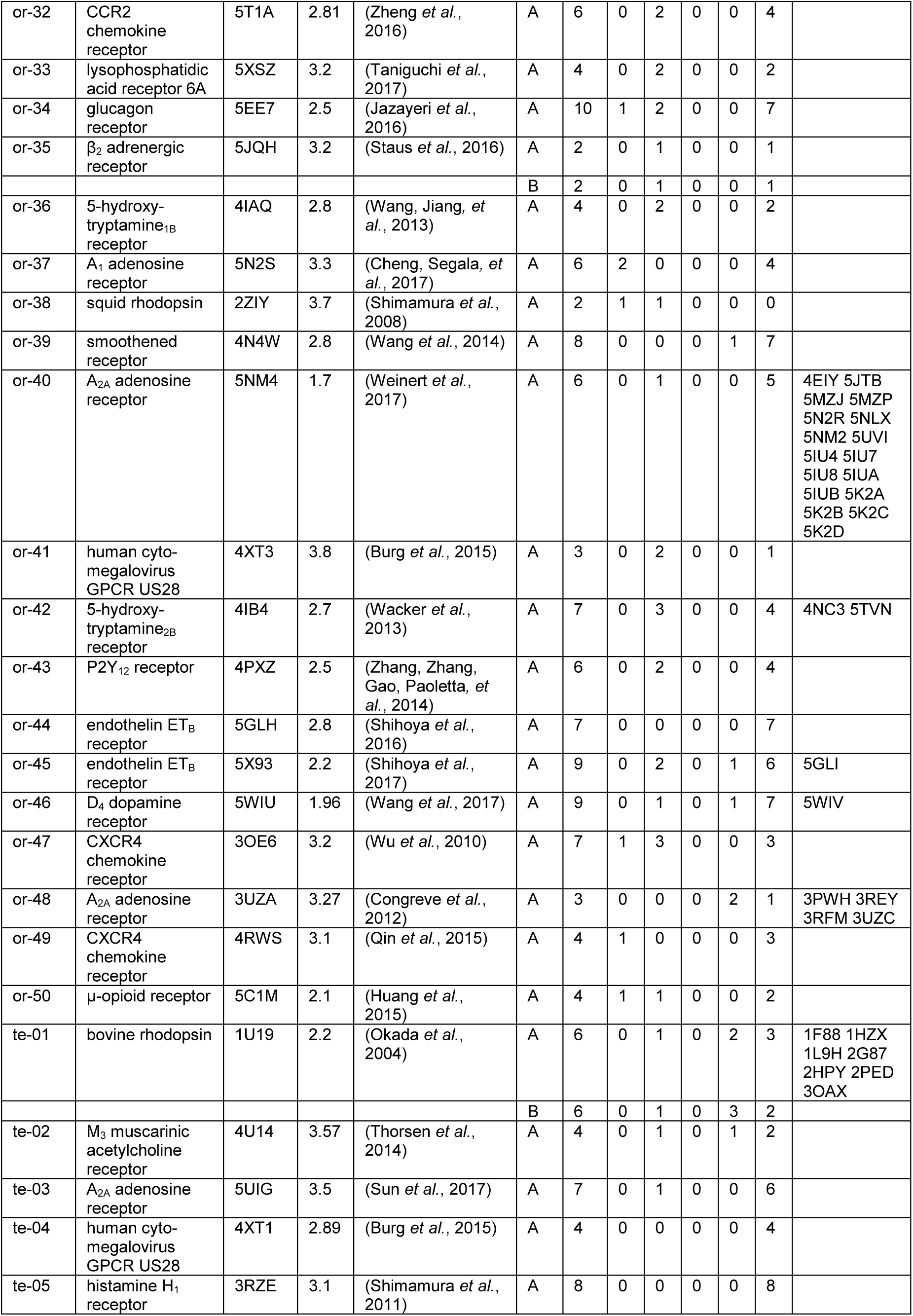
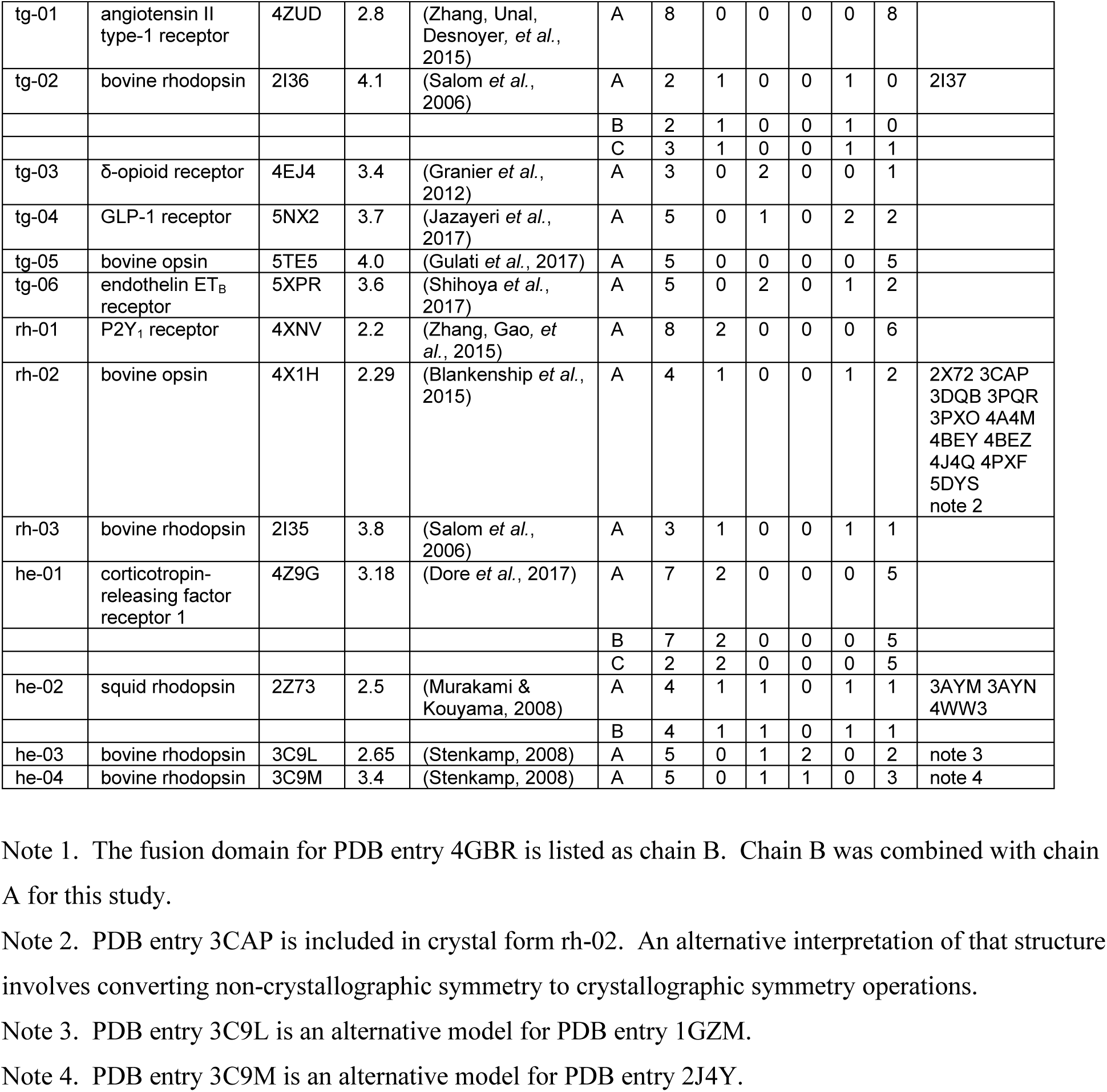
Crystal forms and GPCR-GPCR interactions. The PDB identification code for the exemplar structure is given for each crystal form. The number of packing interfaces is provided for each crystallographically unique GPCR chain in that structure. The numbers of parallel, antiparallel, crisscross and other interfaces as well as the interfaces with bSAS values less than 400 Å^2^ are also listed. The last column contains PDB identification codes for isomorphous structures.

The deposited structural models include many non-protein moieties such as crystallization agents, ligands, agonists, antagonists, post-translational modifications, and peptide or protein inhibitors. Non-covalently bound groups were removed from the structural models since the focus of this study is on protein-protein interactions that might mimic subunit interactions in dimers.

Crystal packing environments for the GPCR molecules were generated and analyzed using a mix of programs in the CCP4 program suite (Winn et al., 2011) and locally written programs. Space group symmetry operations were applied using pdbset to generate the molecules surrounding each GPCR in their crystalline lattice. Molecules were considered part of the packing environment for each crystallographically unique GPCR if they had atoms within 4.0 Å of atoms in the central (or base) molecule. The CCP4 superpose program (Krissinel & Henrick, 2004) was used to align the environments on the basis of the central GPCRs. The packing environments were further dissected to generate pairs of interacting GPCRs. For each pair, buried solvent accessible surfaces (bSAS) were calculated using areaimol (Lee & Richards, 1971, Saff & Kuijlaars, 1997).

XtalView (McRee, 1999) and Coot (Emsley et al., 2010) were used for molecular visualization. Figures were generated using XtalView (McRee, 1999), MOLSCRIPT (Kraulis, 1991) and Raster3d (Merritt & Bacon, 1997).

## 3. Results and Discussion

As of November 1, 2017, models for 215 GPCR crystal structures were available in the PDB. If two structures were of the same GPCR with similar unit cell dimensions and identical space group assignments, they were considered isomorphous. For each set of isomorphous structures, the one with the highest resolution was chosen as the exemplar for that crystal form. The assignment of isomorphous structures was not confirmed by comparison of diffraction patterns.

There are 121 unique crystal forms, some of which contain more than one molecule in the asymmetric unit, and this leads to 173 crystallographically unique GPCR molecules, i.e., GPCRs found in different packing environments. The number of close neighbors for each of those varies from 0 to 12 GPCR molecules. (For some of the structures where a GPCR is co-crystallized with other proteins, e.g., its G protein or a crystallization chaperone, there may be no or few GPCR-GPCR interactions.) A 400 Å^2^ cutoff in buried solvent accessible surface areas (bSAS) per protein molecule limited the pairs of GPCRs for further consideration to 319 (out of 1033 possible) even though the cutoff is significantly smaller than the bSAS values for biologically significant oligomers (Janin, 1997). The bSAS calculations were carried out largely to identify the molecular regions involved in GPCR-GPCR interactions.

The pairs of GPCRs were inspected visually and classified into four groups, parallel dimers, antiparallel dimers, crisscross dimers, and others (see Table 1). 71 pairs contained GPCRs oriented parallel to one another as you would expect for pairs that could possibly interact within a membrane. The exact orientation of these molecules with respect to a membrane is unknown, as is the membrane’s location relative to the long axis of the molecules. The membrane is assumed to be roughly perpendicular to the axes of the transmembrane helices. In that case, the molecules in these “parallel” orientations would have their N-termini on one side of the membrane and their C-termini on the other.

137 of the interfaces involve pairs of receptors with the helices of one GPCR opposite in orientation relative to the other GPCR. Twenty interfaces contain molecules with their transmembrane domains roughly perpendicular to each other. Finally, there are 91 interfaces not involving contacts between the transmembrane domains. Many of the crystal structures are of GPCRs fused with other proteins to enhance their stability and/or crystallization properties. The interactions holding the crystals together in some cases are between the intra- and extra-cellular loops and the fusion domains. Some of these represent modes of interaction that might be found in native membranes, but they have been omitted from this study.

The 71 interfaces with GPCRs in parallel orientations will be called dimers for convenience, but it should be remembered that it is unknown if they form stable complexes in membrane environments. The molecules forming each dimer are listed in Tables 2 and 3 along with information allowing their identification in the PDB. The first molecule listed for a dimer is considered the base or central molecule used for superpositions of the dimers.

**Table 2.**
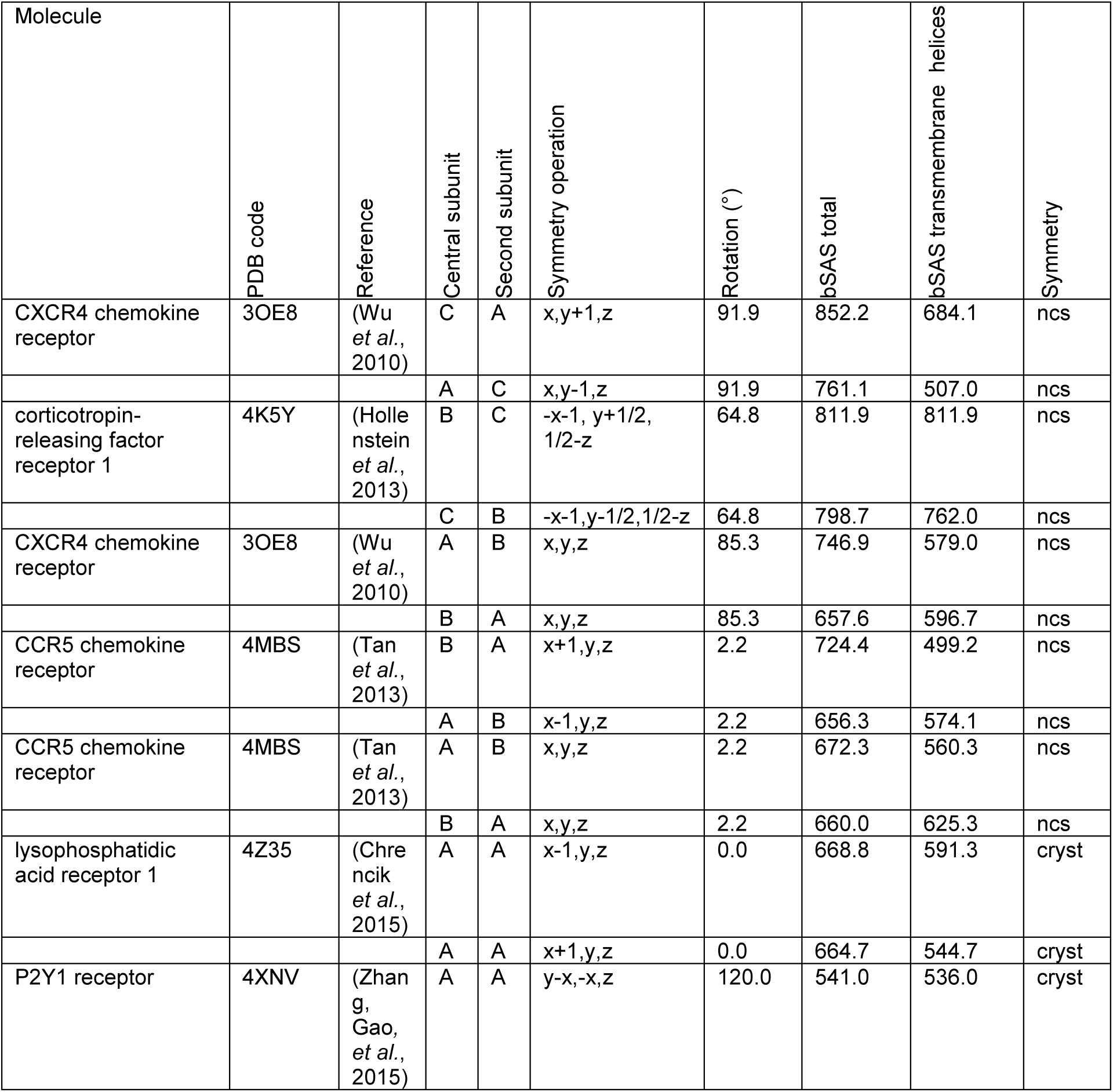
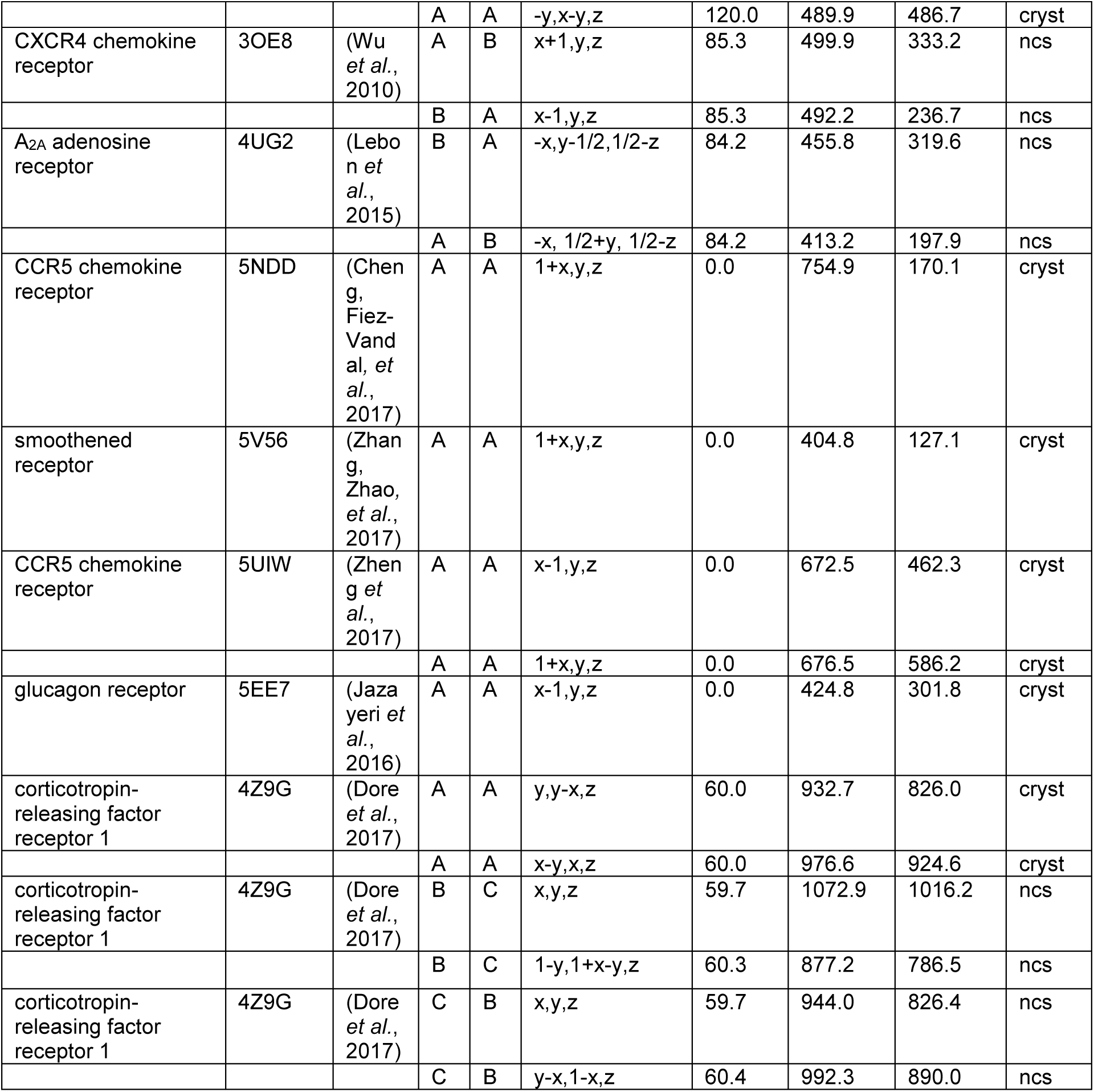
Interfaces between parallel GPCRs not related by two-fold rotational symmetry. The central subunit is positioned as in the deposited PDB file. The symmetry operation is applied to the second chain’s fractional coordinates. The rotation column is derived from the transformation relating the subunits’ transmembrane helices. The buried solvent accessible surface (bSAS) values are for the central molecule and its transmembrane helices. Non-crystallographic (ncs) and crystallographic symmetry (cryst) are noted. Entries with no molecule name listed are the opposing faces for the interfaces listed immediately before them.

| Molecule | PDB code | Reference | Central subunit | Second subunit | Symmetry operation | Rotation (°) | bSAS total | bSAS transmembrane helices | Symmetry |
| --- | --- | --- | --- | --- | --- | --- | --- | --- | --- |
| CXCR4 chemokine receptor | 3OE8 | (Wu *et al.*, 2010) | C | A | x,y+1,z | 91.9 | 852.2 | 684.1 | ncs |
|  |  |  | A | C | x,y-1,z | 91.9 | 761.1 | 507.0 | ncs |
| corticotropin-releasing factor receptor 1 | 4K5Y | (Hollenstein *et al.*, 2013) | B | C | -x-1, y+1/2, 1/2-z | 64.8 | 811.9 | 811.9 | ncs |
|  |  |  | C | B | -x-1,y-1/2,1/2-z | 64.8 | 798.7 | 762.0 | ncs |
| CXCR4 chemokine receptor | 3OE8 | (Wu *et al.*, 2010) | A | B | x,y,z | 85.3 | 746.9 | 579.0 | ncs |
|  |  |  | B | A | x,y,z | 85.3 | 657.6 | 596.7 | ncs |
| CCR5 chemokine receptor | 4MBS | (Tan *et al.*, 2013) | B | A | x+1,y,z | 2.2 | 724.4 | 499.2 | ncs |
|  |  |  | A | B | x-1,y,z | 2.2 | 656.3 | 574.1 | ncs |
| CCR5 chemokine receptor | 4MBS | (Tan *et al.*, 2013) | A | B | x,y,z | 2.2 | 672.3 | 560.3 | ncs |
|  |  |  | B | A | x,y,z | 2.2 | 660.0 | 625.3 | ncs |
| lysophosphatidic acid receptor 1 | 4Z35 | (Chrencik *et al.*, 2015) | A | A | x-1,y,z | 0.0 | 668.8 | 591.3 | cryst |
|  |  |  | A | A | x+1,y,z | 0.0 | 664.7 | 544.7 | cryst |
| P2Y1 receptor | 4XNV | (Zhang, Gao, *et al.*, 2015) | A | A | y-x,-x,z | 120.0 | 541.0 | 536.0 | cryst |
|  |  |  | A | A | -y,x-y,z | 120.0 | 489.9 | 486.7 | cryst |
| CXCR4 chemokine receptor | 3OE8 | (Wu *et al.*, 2010) | A | B | x+1,y,z | 85.3 | 499.9 | 333.2 | ncs |
|  |  |  | B | A | x-1,y,z | 85.3 | 492.2 | 236.7 | ncs |
| A<sub>2A</sub> adenosine receptor | 4UG2 | (Lebo n *et al.*, 2015) | B | A | -x,y-1/2,1/2-z | 84.2 | 455.8 | 319.6 | ncs |
|  |  |  | A | B | -x, 1/2+y, 1/2-z | 84.2 | 413.2 | 197.9 | ncs |
| CCR5 chemokine receptor | 5NDD | (Chen g, Fiez-Vand al, *et al.*, 2017) | A | A | 1+x,y,z | 0.0 | 754.9 | 170.1 | cryst |
| smoothened receptor | 5V56 | (Zhan g, Zhao, *et al.*, 2017) | A | A | 1+x,y,z | 0.0 | 404.8 | 127.1 | cryst |
| CCR5 chemokine receptor | 5UIW | (Zhen g *et al.*, 2017) | A | A | x-1,y,z | 0.0 | 672.5 | 462.3 | cryst |
|  |  |  | A | A | 1+x,y,z | 0.0 | 676.5 | 586.2 | cryst |
| glucagon receptor | 5EE7 | (Jaza yeri *et al.*, 2016) | A | A | x-1,y,z | 0.0 | 424.8 | 301.8 | cryst |
| corticotropin-releasing factor receptor 1 | 4Z9G | (Dore *et al.*, 2017) | A | A | y,y-x,z | 60.0 | 932.7 | 826.0 | cryst |
|  |  |  | A | A | x-y,x,z | 60.0 | 976.6 | 924.6 | cryst |
| corticotropin-releasing factor receptor 1 | 4Z9G | (Dore *et al.*, 2017) | B | C | x,y,z | 59.7 | 1072.9 | 1016.2 | ncs |
|  |  |  | B | C | 1-y,1+x-y,z | 60.3 | 877.2 | 786.5 | ncs |
| corticotropin-releasing factor receptor 1 | 4Z9G | (Dore *et al.*, 2017) | C | B | x,y,z | 59.7 | 944.0 | 826.4 | ncs |
|  |  |  | C | B | y-x,1-x,z | 60.4 | 992.3 | 890.0 | ncs |

**Table 3.**
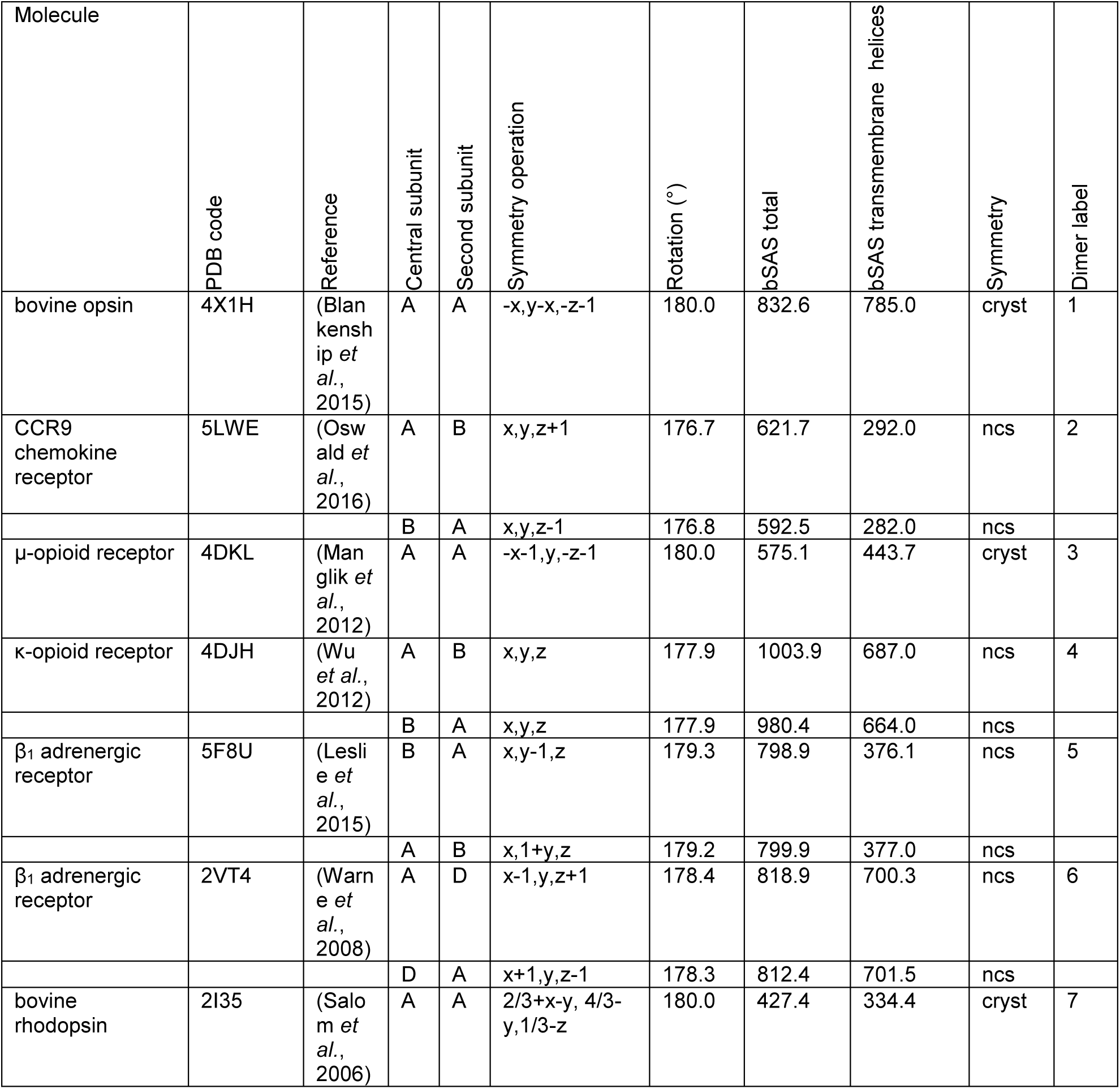
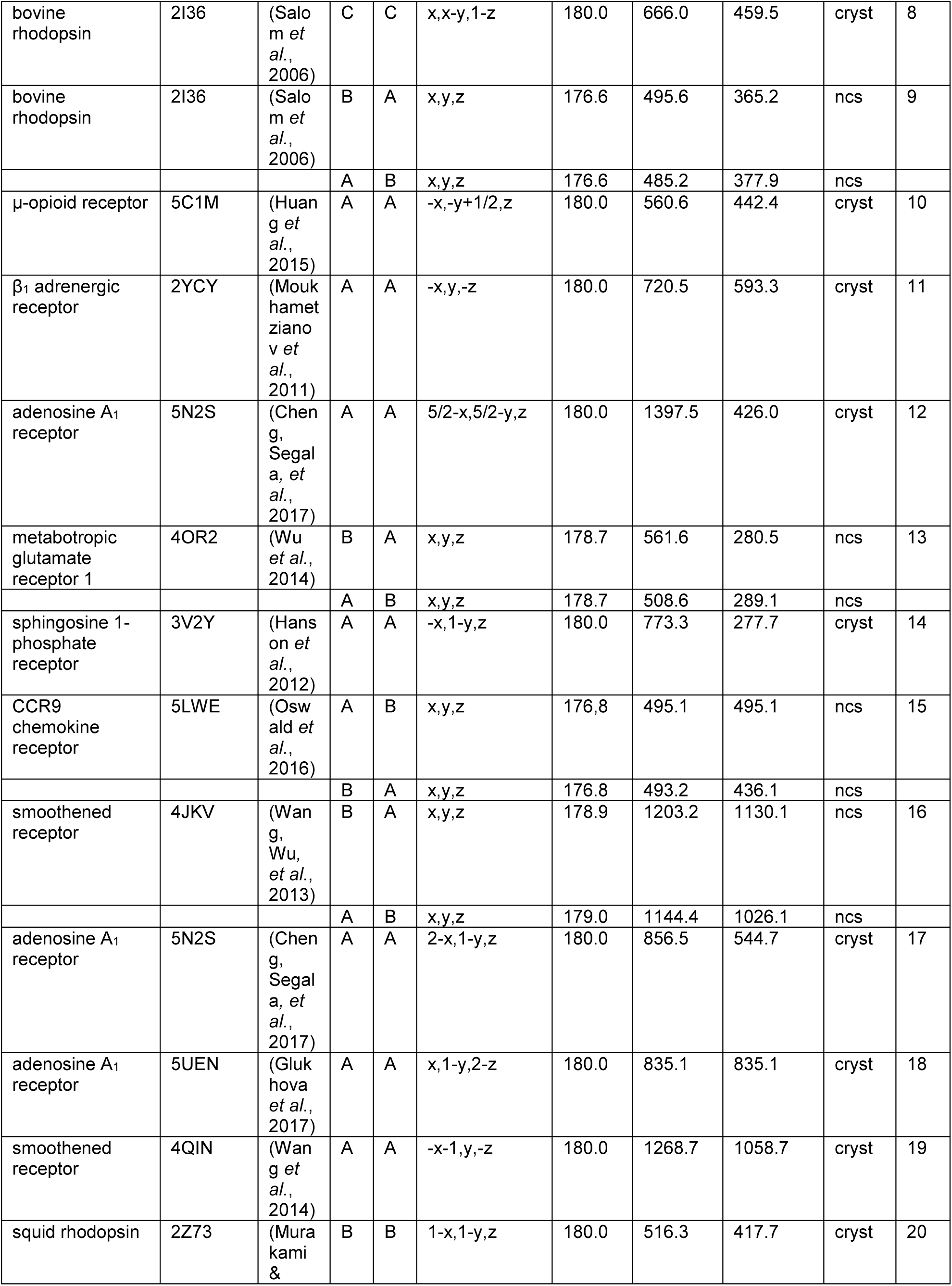
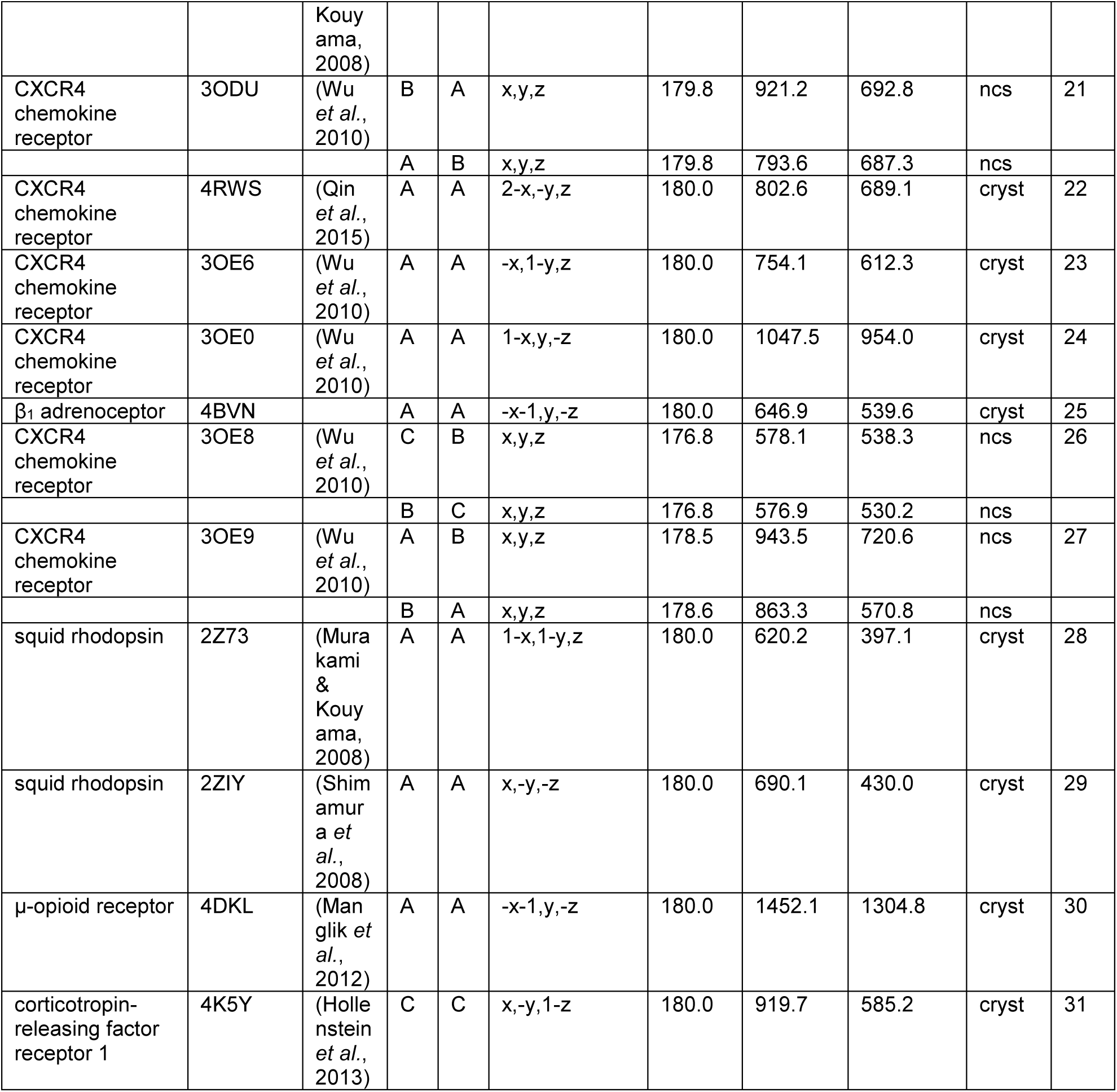
Interfaces between parallel GPCRs related by two-fold rotational symmetry. The central subunit is positioned as in the deposited PDB file. The symmetry operation is applied to the second chain’s fractional coordinates. The rotation column is derived from the transformation relating the subunits’ transmembrane helices. The buried solvent accessible surface (bSAS) values are for the central molecule and its transmembrane helices. Non-crystallographic (ncs) and crystallographic symmetry (cryst) are noted. Entries with no molecule name listed are the opposing faces for the interfaces listed immediately before them. The dimer label column is used throughout the rest of the paper to identify the dimers.

The GPCRs in the parallel dimers are related by combinations of rotations and translations. The 29 interfaces listed in Table 2 involve GPCRs not related by two-fold rotation axes. Eleven of the interfaces involve proteins related by translational symmetry with rotations of 0.0° or 2.2°. Four of the eleven interfaces have two entries in the table because both molecules forming the interface bury more than 400 Å^2^ of bSAS. The other three interfaces have only one entry each in the table because one of the two GPCRs buries less than 400 Å^2^ in the interface. The interactions involved in all eleven are heterologous, and when repeated, lead to infinite oligomers, not discrete dimers. However, these arrays might account for those observed for rhodopsin in native membranes (Liang et al., 2003).

There are 18 parallel interfaces involving molecules related by approximate or exact three-, four-, and six-fold rotational axes perpendicular to the assumed membrane plane. These interactions are also heterologous, so again the interfaces appear twice in Table 2. There are three interfaces between molecules related by approximate six-fold rotation axes, and one with exact six-fold symmetry. Four interfaces have approximate four-fold symmetry, and one has exact three-fold rotational symmetry. The exact rotation axes result in a cyclic hexamer or cyclic trimer. Subunits related by approximate n-fold axes do not form closed cyclic oligomers. Slight conformational changes could result in these approximate symmetry axes becoming to exact rotations, in which case, additional cyclic oligomeric forms might be generated.

The 42 remaining parallel interfaces contain GPCRs with transmembrane domains related by two-fold rotational symmetry axes perpendicular to the membrane plane. Crystallographic two-fold axes relate the GPCRs in 20 of these interfaces, and because the interfaces were generated for each of the crystallographically unique GPCRs, these interfaces appear only once in Table 3.

Twenty-two pairs involve non-crystallographic two-fold axes and appear twice in the table. Strictly speaking, the interactions between the molecules in these dimers are heterologous. The two molecules in a dimer are not crystallographically identical, and the inter-protein interactions are not entirely symmetric. However, most of the angles of rotation relating the pairs of molecules are close to 180° (the largest deviation from 180° being 3.4°). Comparison of the bSAS for the molecules in these dimers show that for eight of the pairs, the difference in bSAS between the two interacting molecules is less than 5% of the larger bSAS value. The molecules in these eight pairs appear to be nearly identical, so the one with the larger bSAS, has been used as the archetype for this dimer. For three of the interfaces (labelled 13, 21, and 27 in Table 3), larger differences in the bSAS for the molecules involved indicate that these interactions are more heterologous. To avoid over-counting of particular structures, only the GPCR with the larger bSAS will be considered further.

Figure 1 displays the result of superposing the 31 dimers listed in Table 3 that contain GPCRs related by two-fold rotation axes. The dimers were oriented by superposing the helices of each base molecule on the transmembrane helices of chain A of PDB entry 4X1H (Blankenship et al., 2015) (opsin). A line connecting the centers of mass of the molecules in each pair was generated (see Figure 1a), and all 31 lines are shown in Figure 1b.

**Figure 1.**
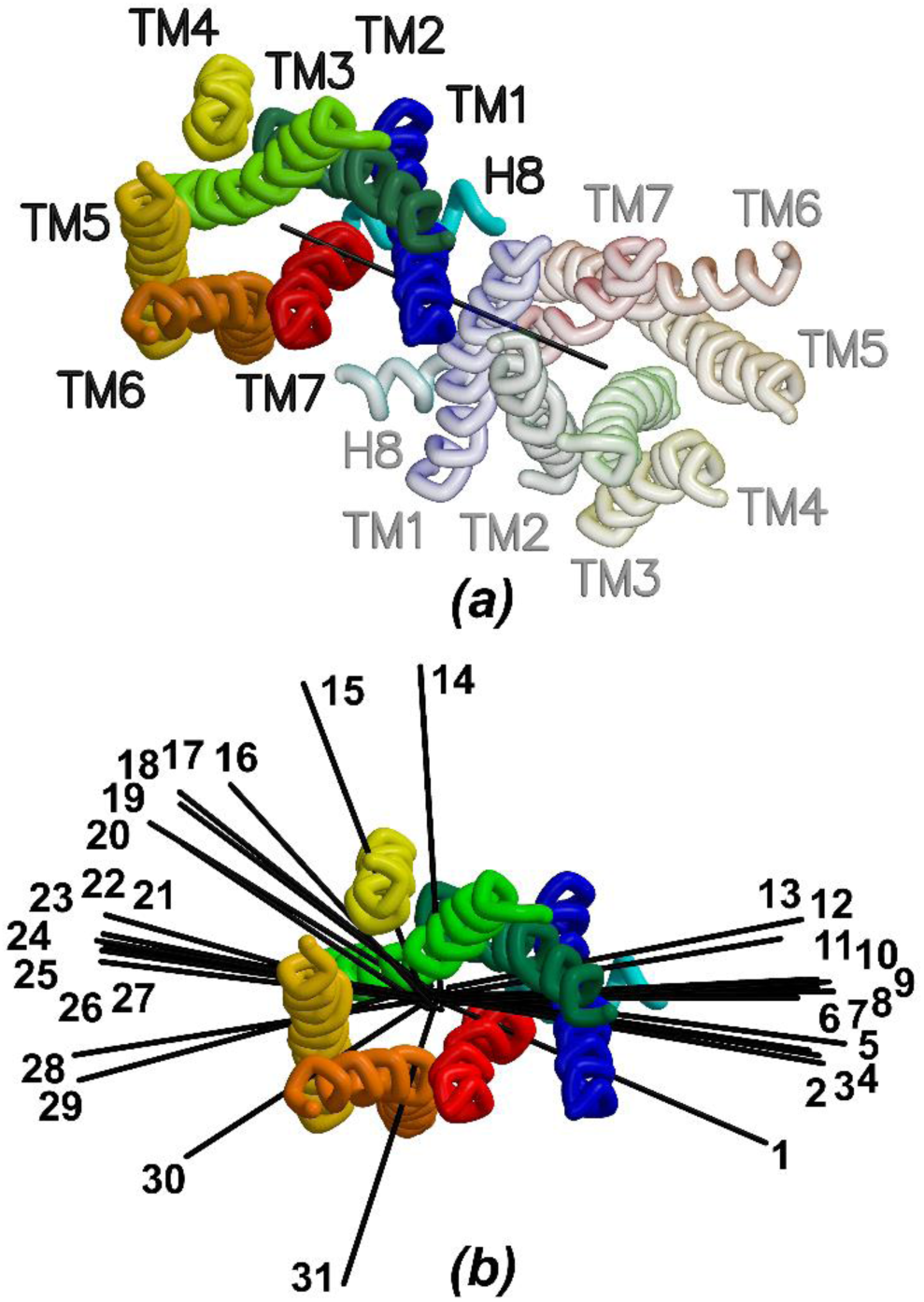
Schematic showing locations of lines-of-center of dimer subunits. (a) Two GPCRs in a dimer with a line joining their centers viewed perpendicular to the plane of the membrane. TM1 shown in blue, TM2 in dark green, TM3 in chartreuse, TM4 in yellow, TM5 in gold, TM6 in orange, TM7 in red, and H8 in cyan. The colors for the second subunit are the same, but in transparent mode. (b) Similar plot, but showing the lines connecting the centers of each of the 31 parallel dimers showing two-fold rotational symmetry (see Table 3).

Dimers 1–13 (see listing in Table 3) form a cluster on the right of Figure 1b and contain receptors interacting mainly through helices TM1, TM2 and H8. Those on the left (14–31) are formed by interactions between helices TM3, TM4, TM5, and TM6. Table 4 lists the helices in the base molecules forming contacts in the dimer interfaces, and Table 5 lists the specific residues in the base molecules involved in each interface.

**Table 4.** Structural elements of the central molecule forming the dimer interface.

| Dimer label<br>(from Table 3) | TM1 | TM2 | TM3 | TM4 | TM5 | TM6 | TM7 | H8 |
| --- | --- | --- | --- | --- | --- | --- | --- | --- |
| 1 | x | x |  |  |  | x | x | x |
| 2 | x |  |  |  |  |  |  |  |
| 3 | x | x |  |  |  |  |  | x |
| 4 | x | x | x |  |  |  |  | x |
| 5 | x | x |  |  |  |  |  | x |
| 6 | x | x |  |  |  |  |  |  |
| 7 | x | x |  |  |  |  |  |  |
| 8 | x | x |  |  |  |  | x | x |
| 9 | x | x |  |  |  |  |  |  |
| 10 | x | x |  |  |  |  |  |  |
| 11 | x | x |  |  |  |  |  |  |
| 12 | x | x |  |  |  |  |  |  |
| 13 | x |  |  |  |  |  |  |  |
| 14 |  |  | x | x |  |  |  |  |
| 15 |  |  |  | x |  |  |  |  |
| 16 |  |  | x | x | x |  |  |  |
| 17 |  |  | x | x | x |  |  |  |
| 18 |  |  | x | x | x |  |  |  |
| 19 |  |  | x | x | x | x |  |  |
| 20 |  |  |  | x | x |  |  |  |
| 21 |  |  |  |  | x | x |  |  |
| 22 |  |  | x | x | x |  |  |  |
| 23 |  |  |  |  | x | x |  |  |
| 24 |  |  | x | x | x | x |  |  |
| 25 |  |  | x | x | x |  |  |  |
| 26 |  |  |  |  | x | x |  |  |
| 27 |  |  | x |  | x | x |  |  |
| 28 |  |  | x |  | x |  |  |  |
| 29 |  |  | x |  | x |  |  |  |
| 30 |  |  |  |  | x | x |  |  |
| 31 |  |  |  |  | x | x | x |  |

**Table 5.** Residues at the dimer interfaces. Generic GPCRdb numbering taken from http://gpcrdb.org/. (a) Dimers 1–13. (b) Dimers 14–20. (c) Dimers 21–31.

| Dimer label (see Table 3) | 1 | 2 | 3 | 4 | 5 | 6 | 7 | 8 | 9 | 10 | 11 | 12 | 13 |
| --- | --- | --- | --- | --- | --- | --- | --- | --- | --- | --- | --- | --- | --- |
| Residue number | bovine opsin | CCR9 chemokine receptor | μ-opioid receptor | κ-opioid receptor | $\beta_1$ adrenergic receptor | $\beta_1$ adrenergic receptor | bovine rhodopsin | bovine rhodopsin | bovine rhodopsin | μ-opioid receptor | $\beta_1$ adrenergic receptor | adenosine A<sub>1</sub> receptor | metabotropic glutamate receptor 1 |
| 1x29 |  |  |  | P56 |  |  |  |  |  | M65 |  |  |  |
| 1x32 |  |  |  | P59 | E41 | E41 |  |  |  |  |  |  |  |
| 1x33 | S38 |  |  | V60 |  | A42 |  |  |  | I69 | A42 |  |  |
| 1x34 | M39 |  |  |  |  |  |  |  |  |  |  |  |  |
| 1x35 |  |  |  |  |  | M44 |  |  |  |  |  |  |  |
| 1x36 |  |  | M72 | T63 | S45 | S45 |  |  |  |  | S45 |  |  |
| 1x37 | A42 | P53 |  | A64 | L46 | L46 |  |  |  |  | L46 |  |  |
| 1x39 |  |  |  | Y66 |  |  |  |  |  |  |  |  |  |
| 1x40 | F45 | W56 | S76 | S67 | A49 | A49 | F45 |  | F45 |  | A49 |  |  |
| 1x41 | L46 | L57 |  | V68 |  | L50 |  |  |  |  | L50 |  |  |
| 1x43 |  |  |  | F70 |  | V52 | I48 |  |  |  | V52 |  |  |
| 1x44 |  | I60 |  | V71 | L53 | L53 | M49 | M49 | M49 |  | L53 | L1126 |  |
| 1x47 |  |  |  |  |  | V56 |  | F52 | F52 |  | V56 | V1129 |  |
| 1x48 |  |  |  |  |  |  |  | P53 | P53 |  |  | P1130 | L605 |
| 1x51 |  |  |  |  |  |  |  |  |  |  |  | V1133 | L608 |
| 1x52 |  |  |  |  |  |  |  |  |  |  |  | L1134 | F609 |
| 1x55 |  |  |  |  |  |  |  |  |  | Y91 |  | W1137 | L612 |
| 1x56 |  |  |  |  |  |  |  |  |  |  |  |  | I613 |
| 1x59 |  |  |  |  |  |  |  |  |  | R95 |  |  | L616 |
| 1x60 |  |  |  |  |  |  |  |  |  |  |  |  | Y617 |
| 2x55 |  |  |  |  |  |  |  |  |  |  |  | A1165 |  |
| 2x58 |  |  |  | P113 |  | P96 |  |  |  |  | P96 |  |  |
| 2x59 |  |  |  | F114 |  | F97 |  |  |  |  |  |  |  |
| 2x61 | L95 |  |  | S116 |  | A99 |  |  |  |  | A99 |  |  |
| 2x62 |  |  | V126 | T117 |  | T100 | Y96 | Y96 | Y96 | V126 | T100 |  |  |
| 2x65 | L99 |  | L129 | L120 |  |  |  |  |  | L129 |  |  |  |
| 2x66 |  |  | M130 |  | R104 | R104 |  | H100 | H100 | M130 | R104 |  |  |
| 3x20 |  |  |  | F126 |  |  |  |  |  |  |  |  |  |
| 6x32 | E249 |  |  |  |  |  |  |  |  |  |  |  |  |
| 6x35 | R252 |  |  |  |  |  |  |  |  |  |  |  |  |
| 7x43 | T297 |  |  |  |  |  |  |  |  |  |  |  |  |
| 7x47 | V300 |  |  |  |  |  |  |  |  |  |  |  |  |
| 7x48 | Y301 |  |  |  |  |  |  |  |  |  |  |  |  |
| 7x51 | V304 |  |  |  |  |  |  |  |  |  |  |  |  |
| 7x52 | I305 |  |  |  |  |  |  |  |  |  |  |  |  |
| 7x55 |  |  |  |  |  |  |  | M308 |  |  |  |  |  |
| 7x56 | M309 |  |  |  |  |  |  |  |  |  |  |  |  |
| 8x48 |  |  |  |  |  |  |  |  | K311 |  |  |  |  |
| 8x51 |  |  |  | K338 | R350 |  |  | R314 | R314 |  |  |  |  |
| 8x54 |  |  | F347 | F341 | F353 |  |  |  |  |  |  |  |  |
| 8x55 |  |  |  | R342 | K354 |  |  |  |  |  |  |  |  |
| 8x58 |  |  | C351 |  |  |  |  | L321 |  |  |  |  |  |
| 8x59 | C322 |  | I352 | F346 |  |  |  | C322 |  |  |  |  |  |

(b).
| Dimer label<br>(see Table 3) | 14 | 15 | 16 | 17 | 18 | 19 | 20 |
| --- | --- | --- | --- | --- | --- | --- | --- |
| Residue number | sphingosine 1-phosphate<sub>1</sub><br>receptor | CCR9 chemokine receptor | smoothened receptor | adenosine A<sub>1</sub> receptor | adenosine A<sub>1</sub> receptor | smoothened receptor | squid rhodopsin |
| 3x23 | A115 |  |  |  |  |  |  |
| 3x27 | L119 |  |  |  |  |  |  |
| 3x44 |  |  | V333 |  |  |  |  |
| 3x48 |  |  | Y337 |  |  |  |  |
| 3x51 |  |  | H340 |  |  | H340 |  |
| 3x55 |  |  |  | K1215 |  |  |  |
| 3x59 |  |  |  |  | R114 |  |  |
| 4x35 |  |  |  |  | M117 |  |  |
| 4x36 |  |  |  |  | V118 |  |  |
| 4x37 |  | K160 |  |  |  |  |  |
| 4x40 | F158 | L163 |  | R1227 |  |  |  |
| 4x41 |  | Y164 |  | R1228 |  |  |  |
| 4x43 | F161 |  |  |  |  | S358 |  |
| 4x44 |  | M167 |  |  |  | Y359 |  |
| 4x45 |  |  | F360 |  |  |  |  |
| 4x47 | S165 | F170 | L362 |  |  | L362 |  |
| 4x48 |  | T171 | L363 |  |  |  |  |
| 4x51 |  | V174 |  |  |  | S366 |  |
| 4x52 |  |  |  |  |  | L367 | W162 |
| 4x54 | L172 |  |  |  |  |  |  |
| 4x55 |  | A178 | V370 | V1242 | V137 | V370 | L165 |
| 4x56 |  |  | L371 |  |  |  | W166 |
| 4x58 |  |  | V373 |  |  |  |  |
| 4x59 |  | I181 | A374 |  | T141 |  |  |
| 4x62 |  |  | A377 | V1249 | F144 |  |  |
| 4x63 |  |  | V378 | G1250 | G145 |  |  |
| 4x64 |  |  |  |  | W146 |  |  |
| 4x68 |  |  |  |  | S150 |  |  |
| 5x30 |  |  |  |  | E153 |  |  |
| 5x31 |  |  |  |  | R154 |  |  |
| 5x34 |  |  |  |  |  |  | S195 |
| 5x35 |  |  |  |  |  |  | T196 |
| 5x38 |  |  | R398 | E1283 | E178 | R398 | S199 |
| 5x39 |  |  | Y399 |  |  | Y399 |  |
| 5x41 |  |  |  |  |  | A401 |  |
| 5x411 |  |  |  | Y1287 | Y182 |  |  |
| 5x42 |  |  | G402 |  | F183 | G402 | C203 |
| 5x421 |  |  | F403 |  |  | F403 |  |
| 5x45 |  |  | A406 |  |  | A406 |  |
| 5x46 |  |  | P407 | V1292 | V187 |  | L207 |
| 5x49 |  |  |  |  |  | L410 |  |
| 5x52 |  |  |  |  |  | I413 |  |
| 5x53 |  |  | V414 |  |  | V414 |  |
| 5x56 |  |  | Y417 |  |  | Y417 |  |
| 5x57 |  |  | F418 |  |  | F418 |  |
| 5x60 |  |  | R421 |  |  | R421 |  |
| 6x56 |  |  |  |  |  | F474 |  |

(c).
| Dimer label<br>(see Table 3) | 21 | 22 | 23 | 24 | 25 | 26 | 27 | 28 | 29 | 30 | 31 |
| --- | --- | --- | --- | --- | --- | --- | --- | --- | --- | --- | --- |
| Residue number | CXCR4 chemokine receptor | CXCR4 chemokine receptor | CXCR4 chemokine receptor | CXCR4 chemokine receptor | $\beta_1$ adrenoceptor | CXCR4 chemokine receptor | CXCR4 chemokine receptor | squid rhodopsin | squid rhodopsin | μ-opioid receptor | corticotropin-releasing factor receptor 1 |
| 3x51 |  | Y135 |  | Y135 | Y140 |  | Y135 | Y134 | Y134 |  |  |
| 3x52 |  |  |  | L136 | L141 |  |  |  |  |  |  |
| 3x55 |  |  | T144 |  |  |  |  |  |  |  |  |
| 3x56 |  |  |  | H140 | S145 |  |  |  |  |  |  |
| 4x35 |  | R146 |  | R146 |  |  |  |  |  |  |  |
| 4x41 |  |  |  |  | R157 |  |  |  |  |  |  |
| 5x32 | N192 |  |  | N192 |  |  | N192 |  |  |  |  |
| 5x33 | D193 |  | D193 | D193 |  | D193 |  |  |  | W226 |  |
| 5x34 | L194 |  | L194 | L194 |  | L194 | L194 |  |  | Y227 |  |
| 5x35 | W195 |  | W195 | W195 |  | W195 | W195 |  |  |  |  |
| 5x37 | V197 | V197 | V197 | V197 |  | V197 | V197 |  |  | N230 |  |
| 5x38 | V198 | V198 | V198 | V198 | A206 | V198 | V198 |  |  | L231 |  |
| 5x41 | F201 | F201 | F201 | F201 | I209 | F201 | F201 |  |  | I234 |  |
| 5x42 | Q202 |  | Q202 |  | A210 | Q202 | Q202 |  |  |  | Y270 |
| 5x44 |  |  |  |  |  |  |  |  |  | F237 |  |
| 5x45 | M205 |  | M205 |  | I213 | M205 | M205 |  |  | I238 |  |
| 5x46 |  |  |  |  | I214 |  |  |  |  |  |  |
| 5x48 |  |  |  |  |  |  |  |  |  | I242 |  |
| 5x49 | L210 | L210 | L210 | L210 | I218 | L210 |  |  |  |  |  |
| 5x52 |  |  |  |  |  |  | I213 |  | L214 | L246 |  |
| 5x55 |  |  |  |  |  |  |  |  |  | T249 |  |
| 5x56 |  |  |  |  |  |  |  | F218 | F218 |  |  |
| 5x58 |  |  |  |  |  |  |  |  |  | Y252 |  |
| 5x59 |  |  |  |  |  |  |  | F221 | F221 | G253 |  |
| 5x60 |  |  |  |  |  |  |  | N222 | N222 |  |  |
| 5x61 |  |  |  |  |  |  |  |  |  | M255 |  |
| 5x62 |  |  |  |  |  |  |  |  |  | I256 |  |
| 5x63 |  | S224 |  |  | R232 |  |  | M225 | M225 | L257 |  |
| 5x64 |  |  |  | K225 | E233 |  |  |  |  |  |  |
| 5x66 |  |  |  |  |  |  |  |  | S228 |  |  |
| 5x67 |  | H228 |  | H228 |  |  |  | N229 | N229 |  |  |
| 5x70 |  |  |  |  |  |  |  |  | K232 |  |  |
| 6x35 |  |  |  |  |  |  |  |  |  | R280 |  |
| 6x38 |  |  |  |  |  |  |  |  |  | L283 |  |
| 6x42 |  |  |  |  |  |  |  |  |  |  | P321 |
| 6x46 |  |  |  |  |  |  |  |  |  |  | I325 |
| 6x49 |  |  |  |  |  |  |  |  |  | T294 | M328 |
| 6x50 |  |  |  |  |  |  |  |  |  |  | L329 |
| 6x51 |  |  |  |  |  |  |  |  |  |  | A330 |
| 6x53 |  |  |  |  |  |  |  |  |  | I298 | V332 |
| 6x57 |  |  |  |  |  |  |  |  |  | I302 |  |
| 6x60 |  |  |  |  |  |  |  |  |  | L305 |  |
| 6x61 |  |  |  |  |  |  |  |  |  | I306 |  |
| 6x62 | L266 |  | L266 | L266 |  | L266 | L266 |  |  |  |  |
| 6x63 | L267 |  | L267 | L267 |  | L267 | L267 |  |  |  |  |
| 7x27 |  |  |  |  |  |  |  |  |  |  | E338 |
| 7x33 |  |  |  |  |  |  |  |  |  |  | F344 |

Identification of these general inter-helical interactions is not a new observation. Katritch, Cherezov and Stevens commented on several crystallographic dimers (Katritch et al., 2013), and Baltoumas, Theodoropoulou and Hamodrakas included 16 crystallographic and computationally modeled dimers in their molecular dynamics study (Baltoumas et al., 2016). The systematic search described here provides additional crystallographic examples of possible dimers. This larger set of examples shows the range of structures in the two clusters of dimers.

While the subunits in dimers 1–13 mainly interact via helices TM1, TM2 and H8, the relative orientation of the two-fold rotation axes relating the subunits in these dimers varies across the set. The subunits in dimers 1, 2, 3, 4, and 5 are arranged such that their two-fold axes are parallel to the long axes of the central receptor shown in Figure 1. Figure 2a is a view of dimer 4, looking approximately down its two-fold rotation axis and is representative of this group of dimers. In this group, H8 from the left subunit is above that from the right subunit, see Figure 2b.

**Figure 2.**
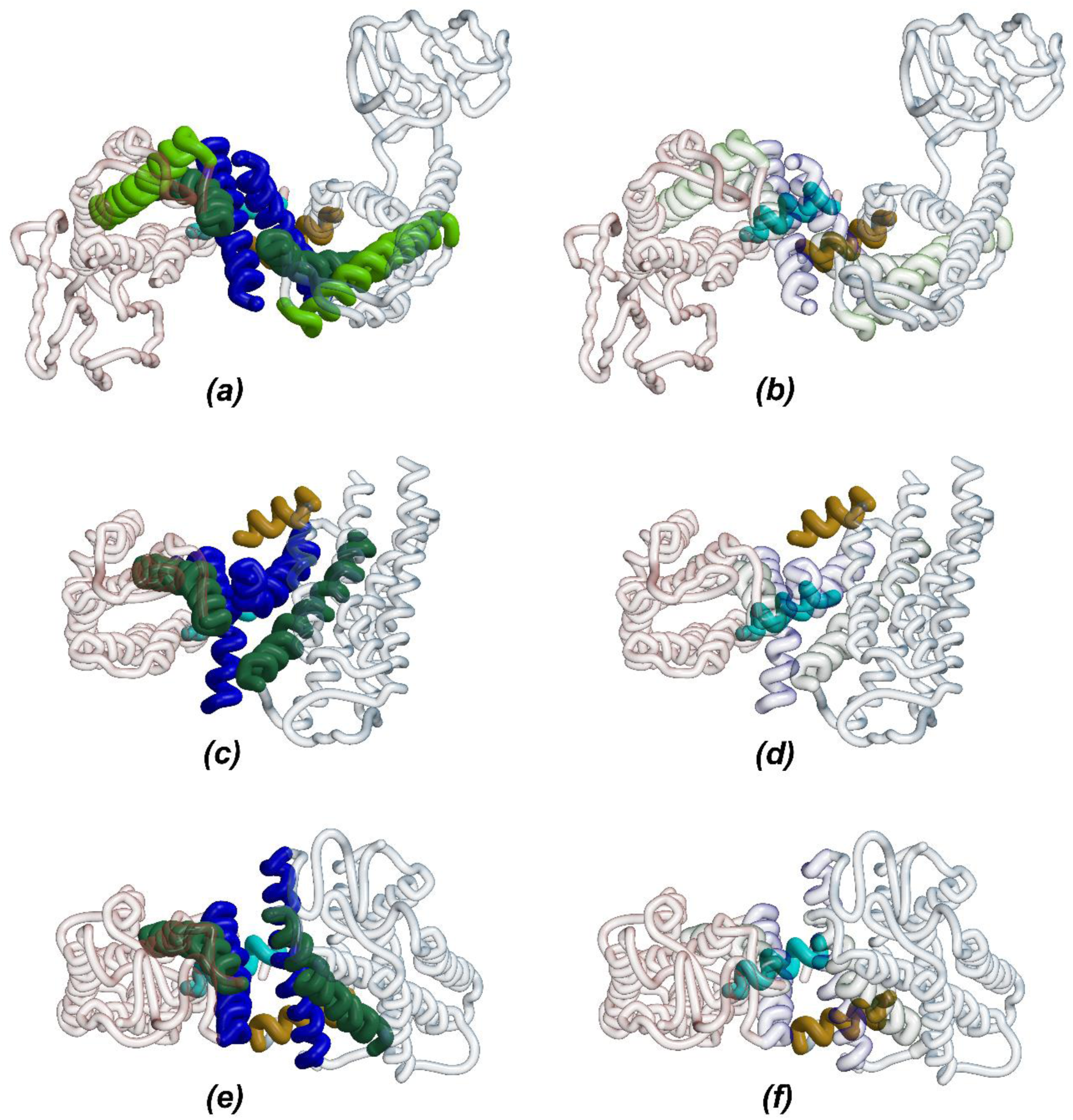
Selected dimers formed using TM1-3 at their interfaces. The central and second molecules are shown as transparent red or blue alpha carbon tracings. Transmembrane helices at the interface are shown in bold colors. TM1 shown in blue, TM2 in dark green, TM3 in chartreuse. H8 from the central and second molecules shown in cyan and brown. (a) Dimer 4 looking down on the extracellular side of the GPCRs. (b) Same view of dimer 4, but clipped to show H8. (c) Dimer 6. (d) Same view as in (b). (e) Dimer 7. (f) Same view as in (b).

Dimers 6, 10, and 11 form another group with their two subunits twisted about the line between their centers when compared to the dimers in the first group (compare Figure 2c and 2a). As seen for dimer 6, the positions of H8 switch such that H8 from the left subunit is now below that from the right (Figure 2d). This structural difference positions the opposite side of the helix for subunit-subunit interactions compared to the dimers represented in Figure 2a.

A third set of dimers (7, 8, and 9) also contain subunits twisted about their line of centers relative to the first group (see Figure 2e), however, the sense of this twist is opposite to that of the second group. Associated with this different twist, H8 from the left subunit is again above that from the right (see Figure 2f). In dimers 12 and 13, there are no H8 helices, and this relaxes any restraint the helix places on subunit-subunit orientation. The connection between the orientation of the subunits and the position of H8 in the interface was made by Baltoumas, *et al*. (Baltoumas et al., 2016), and the role of H8 in GPCR functioning has been noted (Sato et al., 2016) with a focus on its interactions with other structural elements of its GPCR and on interactions between the GPCR and G proteins. It is also possible that amino acid sequence differences or physical conditions controlling helix folding of H8 might modulate signaling by affecting the role of H8 in the dimer interface.

On the other side of Figure 1b, the interfaces for dimers 14–31 are formed by inter-helical interactions involving TM3, TM4, TM5, and/or TM6 (see Figure 3 and Table 4). These structures cover a broad range of orientations, but they can also be grouped according to the locations of the two-fold rotation axes relating their subunits. The transmembrane helices in the subunits for these dimers are all roughly perpendicular to the membrane plane, but the rotation axes are found at different positions around the surface of the central molecule. This results in a systematic variation in the helices forming the dimer interactions (see Figure 3). Dimer 18 is representative of dimers 16–20 (see Figure 3a) where TM3, TM4 and TM5 are found at its interface. Dimer 24 is near the center of dimers 21–27 (Figure 3b) where TM6 joins the set of interacting helices. Finally, for dimers 28 (Figure 3c), the dimeric interface has moved along the surface to the point where neither TM4 nor TM6 makes contacts between the subunits. As is the case for dimers 1–13, considerable structural variation is found in interfaces for dimers 14–31, enough to provide several possible beginning points for homology modeling projects.

**Figure 3.**
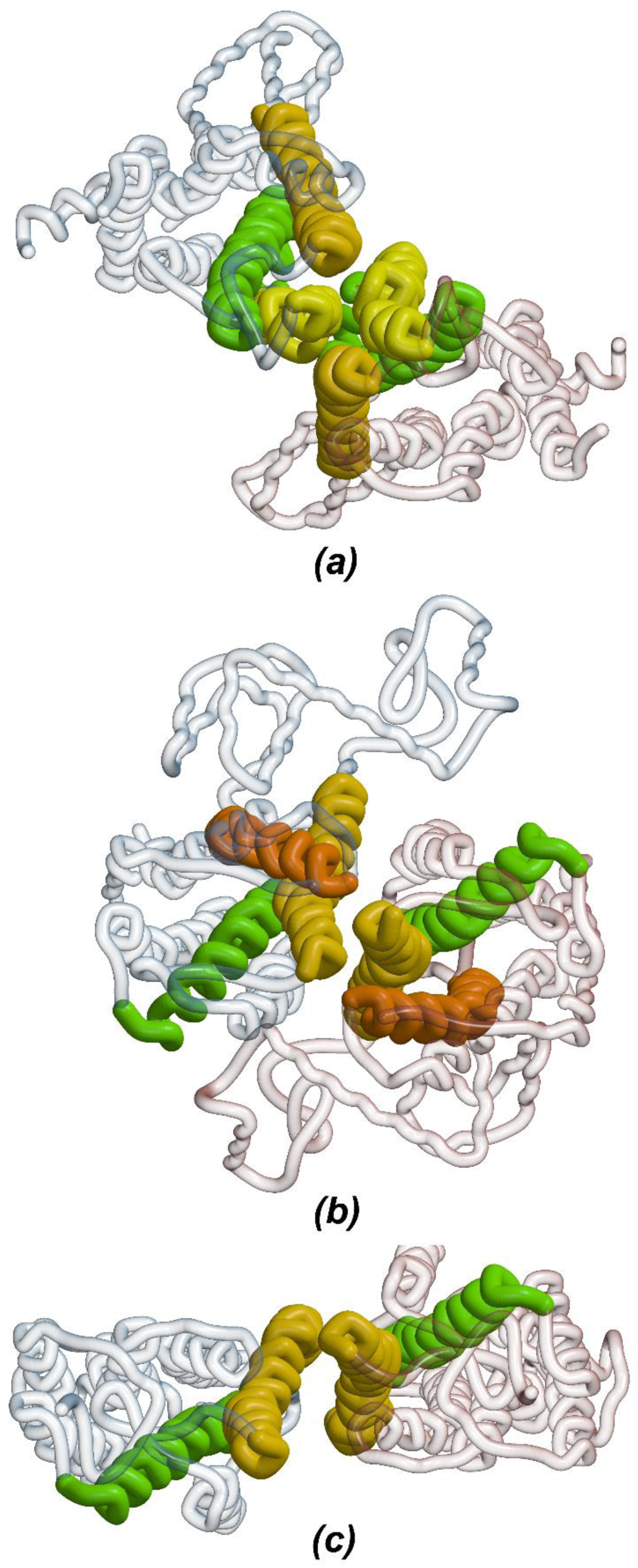
Selected dimers formed using TM3-6 at their interfaces. The central molecule is shown as a transparent red alpha carbon tracing. The second molecule is shown in transparent blue. Transmembrane helices at the interface are shown in bold colors. TM3 in chartreuse, TM4 in yellow, TM5 in gold, TM6 in orange. (a) Dimer 18 looking down on the extracellular side of the GPCRs. (b) Dimer 24. (c) Dimer 28.

The residues in the transmembrane helices in the dimer interfaces are listed in Table 5. As expected for interactions between transmembrane helices, most of the residues involved are hydrophobic. At the ends of the helices, hydrophilic residues can form hydrogen bonds with other polar groups, and a few polar residues are also found in the central parts of the helices. Identification of residue positions that might be crucial for inter-subunit interactions is complicated by multiple occurrences of particular GPCRs in Table 5. For example, while residue position 1x40 (GPCRdb numbering (Isberg et al., 2016)) is located in the interfaces of nine dimers, if β_1_ adrenergic receptor and rhodopsin are counted only once, that number drops to five. Similarly, counting of the CXCR4 dimers only once significantly changes the number of times residues such as 3×51 or 5×38 are found at dimer interfaces.

While the orientation of the GPCRs in the interfaces reported here is consistent with them being in natural membranes, it is not clear that the observed inter-GPCR interactions are strong enough to be biologically relevant. Contributions of “core” residues in stabilizing inter-molecular contacts are major considerations in the eppic webserver (http://www.eppic-web.org/ewui/#) (Duarte et al., 2012) designed to determine oligomeric states of membrane proteins. Most of the dimers listed in Table 3 have small bSAS, and there are few, if any, “core” residues in their interfaces. eppic suggests that the interfaces found in the 31 possible dimers are characteristic of crystal packing interfaces and not biologically relevant ones. Post-translational modifications and bound ligands further complicate this issue since they can stabilize the dimer interactions. Views of the interactions of these moieties with GPCRs are difficult to find since only a few of the low or moderate resolution crystal structure models include these groups.

While none of the entries in Table 3 appear to involve extensive biologically relevant interfaces, it is noteworthy that CXCR4 appears in the table six times (dimers 21–24, 26, 27). As noted in the original structure reports (Wu et al., 2010, Qin et al., 2015), a CXCR4 dimer is found in six different crystal packing arrangements, i.e., six different crystal forms. The six CXCR4 entries in Table 3 form a tight cluster in Figure 1b, and the structures are similar (see Figure 3b). These dimers make use of TM3, TM4, TM5, and TM6 in their inter-subunit interfaces.

Another receptor that appears several times in Table 3 is the β_1_-adrenergic receptor (Huang et al., 2013, Warne et al., 2008, Moukhametzianov et al., 2011, Miller-Gallacher et al., 2014). In dimers 5, 6, and 11, TM1 and TM2 form the major interactions between the subunits. Dimers 6 and 11 are similar, but their subunits are oriented differently from those in dimer 5 (see Figure 2). β_1_-adrenergic receptor dimers can also be formed by interactions using TM3, TM4 and TM5 (dimer 25). As opposed to the case of CXCR4 where the same interaction mode between GPCRs is seen in several crystal packing arrangements, fundamentally different dimers of β_1_-adrenergic receptor need to be considered for studies of its oligomerization.

Rhodopsin/opsin dimers appear four times in Table 3 (numbers 1, 7, 8, and 9). Dimer 1 is formed by subunits that are nearly aligned. Dimers 7, 8, and 9 are similar in structure with subunits twisted relative to one another (Figure 2e). TM1 and TM2 form part of the subunit interfaces in all four dimers. Residues from TM6, TM7 and H8 are found at the interface in dimer 1, while dimer 8 also makes use of TM7 and H8 in its dimer interface. Low resolution electron microscopic, computational (PDB entry 1N3M), crosslinking, and mutagenesis studies (Liang et al., 2003, Jastrzebska et al., 2011, Jastrzebska et al., 2016) resulted in the development and support of a rhodopsin dimer (PDB entry 1N3M) that utilizes TM4 and TM5 in the dimer interface. While this type of dimer is not consistent with the pairs seen in the rhodopsin/opsin crystal structures, it is like dimers 17–25.

## 4. Conclusions

Two major types of GPCR dimers deduced from their crystal structures have been identified by others (Katritch et al., 2013, Baltoumas et al., 2016). Previous studies have focused on particular receptors and their possible oligomers. What is unclear from the published records is whether all packing interactions in known GPCR crystal structures have been considered in those studies. The larger number of experimentally determined GPCR crystal structures available now provides an opportunity to see how varied the members of those clusters are.

In the case of the dimers formed by interactions involving TM1 and TM2, twisting of the subunits around the line between their centers affects the overall relative orientations of the subunits. In addition, this has a major effect on H8 and how it interacts with other parts of its own GPCR and/or its dimeric mate. These structural differences might be associated with the effects of H8 on GPCR function.

Because of the recurring CXCR4 dimers and the rhodopsin modeling, the TM3-6 interface has been the focus of much attention. However, not all GPCR dimers based on this interface make use of it in the same way. There is a continuum of structures using portions of this interface because it covers such a large part of the protein’s transmembrane surface. The variation can be seen in Table 4 where dimers 14–18 mainly have TM3-5 between the subunits, dimers 19–29 use TM3-6, and finally dimers 30-31 are stabilized by interactions of TM5 and TM6. The different pieces of this surface used in forming dimers is a complication of some importance because it expands the number of possible beginning models for computational studies of GPCR dimerization.

Examination of all of the dimeric crystal packing interactions also shows that only certain parts of the transmembrane surfaces of GPCRs are involved in forming parallel dimers. This appears as the systematic split in the dimers in Figure 1b. It is apparent from comparing interfaces from the crystal structures that not all combinations of transmembrane helices can lead to stable dimers. This restriction on which helices form dimeric interfaces is a powerful restraint to be considered in modeling GPCR dimers.

## Acknowledgements

I’m grateful to David Salom, Beata Jastrzebska, Philip Kiser, Isolde Le Trong, David Lodowski, Kris Palczewski, Peter Brzovic, and Dave Teller for encouragement and discussions of this and other GPCR-related topics and to Jeff Godden for technical assistance with the figures.

